# Interspecies generation of functional muscle stem cells

**DOI:** 10.1101/2023.04.12.536533

**Authors:** Seraina A. Domenig, Ajda Lenardič, Joel Zvick, Monika Tarnowska-Sengül, Nicola Bundschuh, Giada Bacchin, Adhideb Ghosh, Ori Bar-Nur

## Abstract

Satellite cells, the stem cells of skeletal muscle tissue, hold a prodigious regeneration capacity. However, low satellite cell yield from autologous or donor-derived muscles precludes adoption of satellite cell transplantation for the treatment of muscle diseases including Duchenne muscular dystrophy (DMD). To address this limitation, here we investigated whether sufficient quantity of satellite cells can be produced in allogeneic or xenogeneic animal hosts. First, we report on exclusive satellite cell production in intraspecies mouse chimeras by injection of CRISPR/Cas9-corrected DMD-induced pluripotent stem cells (iPSCs) into blastocysts carrying an ablation system of host Pax7+ satellite cells. Additionally, injection of genetically-corrected DMD-iPSCs into rat blastocysts produced interspecies rat-mouse chimeras harboring mouse muscle stem cells that efficiently restored dystrophin expression in DMD mice. This study thus provides a proof-of-principle for the generation of therapeutically-competent stem cells between divergent species, raising the possibility of procuring human stem cells in large animals for regenerative medicine purposes.

## Main Text

Muscle degeneration denotes the loss of skeletal muscle mass as a consequence of pathological affliction in the form of sarcopenia, cachexia or muscular dystrophies(1). Following muscle insult, quiescent satellite cells orchestrate a myogenic regeneration program by means of activation and differentiation into transit-amplifying myoblasts that further differentiate into fusion-competent myocytes that merge with damaged multinucleated muscle fibers for tissue repair(1, 2). This step-wise differentiation process is characterized by upregulation of specific transcription factors including paired-box 7 (*Pax7*) in satellite cells and myogenic differentiation 1 (*MyoD*) in myoblasts(1, 2).

DMD is the most common and currently uncurable muscular dystrophy, which arises due to a mutation in the dystrophin gene, a large structural protein that connects skeletal muscle fibers to the extracellular matrix(3-5). In DMD patients, lack of dystrophin renders muscle fibers highly susceptible to breakage due to muscle contraction forces, resulting in increased regeneration cycles by satellite cells(5). However, continuous erosion of myofibers gradually exhausts the regeneration capacity of satellite cells, precipitating muscle fiber replacement with fibrotic and adipogenic tissues over time(6). As a consequence of skeletal muscle wasting, DMD patients become wheelchair-dependent during childhood and eventually succumb to untimely death due to cardiorespiratory failures in the second or third decade of life(6).

A variety of therapeutic interventions are currently being explored for their capacity to restore dystrophin expression(7). Such efforts include gene therapy using overexpression of micro-dystrophin or correction of the DMD mutation by CRISPR/Cas9, typically via use of adeno associated viruses (AAVs)(7). While promising, these approaches still carry various concerns including AAV toxicity, genomic integration or DNA breakage as well as an unfavorable immunological response against repeated AAV treatment or Cas9(8-11). Alternatively, cell-based therapies have been extensively explored for their potential to restore dystrophin expression in DMD animal models via injection of myogenic stem or progenitor cells into dystrophic muscles(12, 13). Such trials aim to add healthy myonuclei to dystrophic myofibers via cell fusion for dystrophin restoration(14). Early endeavors performed in the 1990’s utilizing healthy myoblasts to restore dystrophin expression in DMD patients were unsuccessful, albeit more recent trials reported a better outcome(15-17). One explanation for this unfavorable result is that myoblasts lose *in vivo* engraftment capabilities following extensive *in vitro* expansion(18). As such, major efforts have been directed towards finding means to augment the engraftment potential of myoblasts, or seek additional expandable myogenic cell types that can efficiently restore dystrophin expression *in vivo* following intramuscular injection in DMD animal models(12, 13). Several notable examples include iPSC-derived myogenic precursor cells, teratoma-derived muscle stem cells or directly reprogrammed induced myogenic progenitor cells (iMPCs)(19-25). However, satellite cells are still widely-accepted as the most potent source capable of restoring dystrophin expression, since low number of satellite cells can efficiently engraft and regenerate muscles *in vivo*(18, 26-28). In respect to treating DMD patients, harvesting sufficient number of satellite cells from autologous or donor-derived muscles represents a major challenge for cell-based therapy(12).

Blastocyst complementation is an innovative technology that enables the creation of specific organs, tissues, or cell-types from donor-derived PSCs(29). To this end, PSCs such as embryonic stem cells (ESCs) or iPSCs are injected into blastocysts that carry genetic mutations that impede the formation of specific organs or cell types in animal chimeras, thereby enabling exclusive generation from injected PSCs(29). In recent years, this approach has been utilized to produce cells and organs in intraspecies mouse-mouse or pig-pig chimeras(29). Most notably, this technique was also utilized in an interspecies manner, demonstrating production of organs or cell types in xenogeneic hosts such as pancreas, blood vasculature, kidneys, thymi or germ cells in mice or rats(30-37). However, production of genetically-corrected interspecies adult solid tissue stem cells between different animal species has not been reported to date(29). Here, we set out to combine cellular reprogramming, genome engineering and *in vivo* differentiation of PSCs in mouse-mouse and rat-mouse chimeras to generate allogeneic and genetically-corrected mouse stem cells that can be exploited to restore dystrophin expression in DMD mice.

## Results

### Substantial production of mESC-derived satellite cells in intraspecies chimeras

We commenced our study by setting out to explore whether mESCs can solely produce satellite cells in intraspecies chimeras generated using mouse blastocysts carrying *Pax7-CreERT2* and *Rosa26-loxSTOPlox-Diphteria toxin A* (*R26-LSL-DTA*) homozygous alleles(38, 39). As satellite cells uniquely express the *Pax7* gene in skeletal muscles(40), this system ensures specific ablation of host satellite cells following tamoxifen injection, and can potentially provide a vacant niche receptive for mESC-derived satellite cell colonization in skeletal muscles of chimeras (Figure 1A). To investigate this question, we used lentiviral-transduced Red Fluorescent Protein positive (RFP^+^) KH2-mESCs, which have been previously reported to contribute robustly to mouse chimerism (Figure S1A)(32, 41). Of note, prior to blastocyst injections, RFP^+^mESCs were cultured for 5 days in ‘enhanced’ culture medium to increase chimeric contribution(42). Altogether, we performed three blastocyst injection rounds which produced 28 out of 58 (48%) chimera offspring based on genotyping for the RFP allele and presence of agouti coat color emanating from KH2-mESCs (Figure 1B, 1C and S1B). Furthermore, all mice carried the *R26-LSL-DTA* allele as expected (Figure S1B). Next, we wished to assess whether we can exploit the genetic system to ablate host satellite cells in newborn pups, aiming to create this way a vacant niche for reconstitution with mESC-derived satellite cells during postnatal growth. To this end, we performed tamoxifen injections in 3 day-old chimeric or non-chimeric pups for three consecutive days. This early developmental time point was chosen as it is characterized by rapid muscle growth associated with high proliferation rate of endogenous PAX7^+^ satellite cells(43). Over a course of three weeks after birth, we observed significant bodyweight reduction in tamoxifen injected non-chimeric *Pax7-CreERT2: R26-LSL-DTA* animals, however not in non-injected control animals (Figures 1C and 1D). Notably, intraspecies *Pax7-CreERT2: R26-LSL-DTA /* RFP^+^KH2-mESC chimeras showed no appreciable reduction in size or bodyweight, even when subjected to tamoxifen injection, suggesting rescue by mESCs (Figure 1C and 1D). To validate efficient satellite cell ablation, we harvested leg muscles from non-chimeric *Pax7-CreERT2: R26-LSL-DTA* mice subjected to tamoxifen injections and non-injected controls. We solely detected *Pax*7 expressing satellite cells in non-injected muscle sections, however not in muscles of injected animals (Figure 1E). Next, we observed that RFP^+^KH2-mESCs extensively contributed to skeletal muscle tissue in chimeras as muscle sections exhibited strong RFP expression in resident muscle cells, independent of host satellite cell ablation (Figure 1F). We then assessed whether all PAX7^+^ satellite cells expressed the RFP label in these muscle sections. Unexpectedly, we detected PAX7^+^ satellite cells that were RFP negative, suggesting that either host satellite cells remained following tamoxifen injection, or that transgene silencing occurred in mESC-derived satellite cells (Figure 1G).

**Fig. 1.**
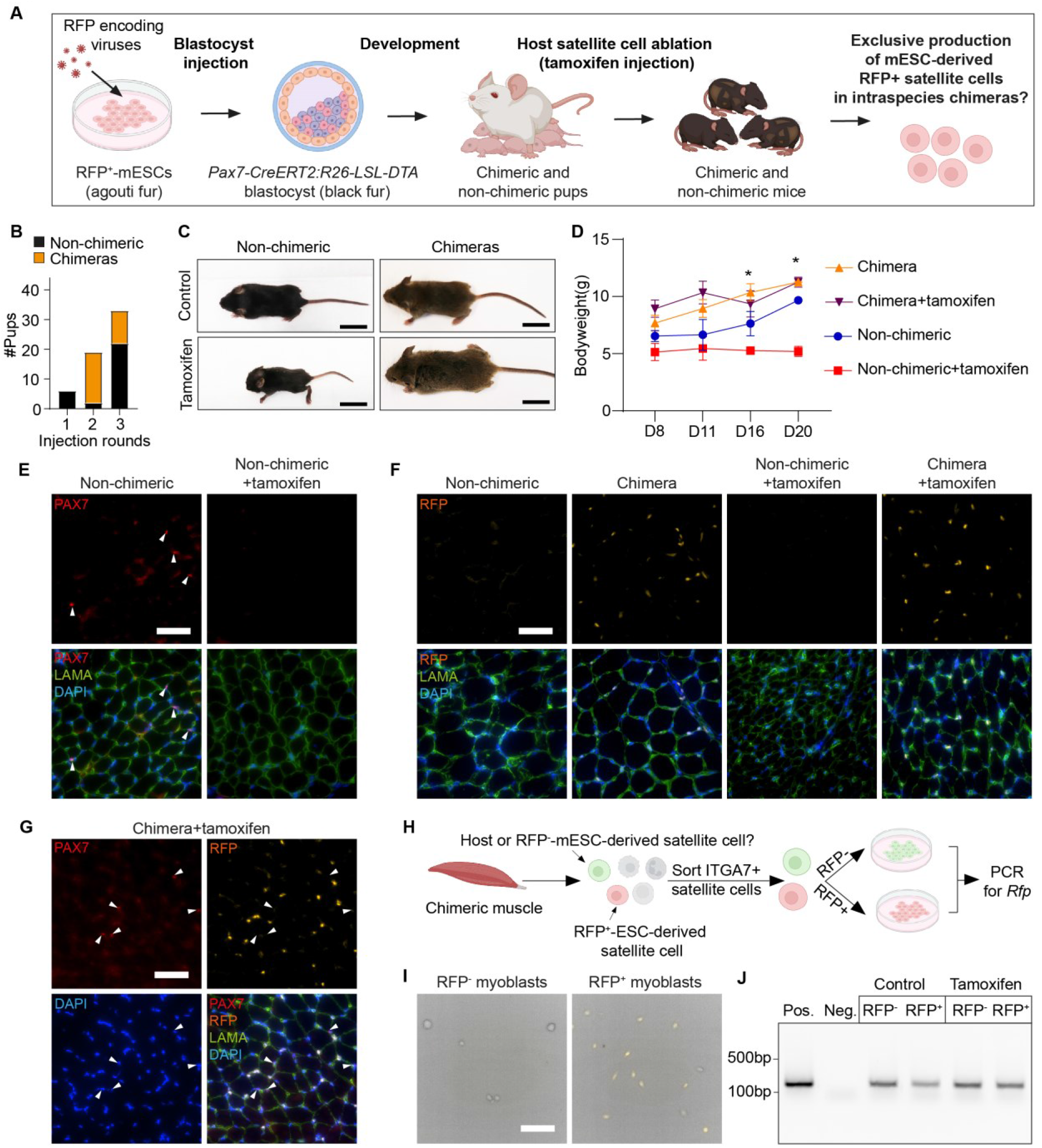
Satellite cells generated in intraspecies chimeras solely from mESCs. **(A)** A schematic of experimental design. RFP, red fluorescent protein; mESCs, mouse embryonic stem cells. **(B)** A graph showing chimera numbers. **(C)** Photos depicting the indicated mouse strains at day 17. Chimerism is represented by agouti coat color. Scale bar, 1cm. **(D)** Graph showing weight during postnatal growth of the indicated strains. Only the non-chimeric+tamoxifen group showed a significant difference in bodyweight compared to the other groups. N≥3, error bars denote SD. Statistical analysis was performed using 2-way ANOVA. ∗p ≤ 0.05. **(E)** Immunofluorescence images of the indicated markers and strains in skeletal muscle cross-sections at day 17. Scale bar, 50µm. **(F)** Immunofluorescence staining for the indicated markers and strains in skeletal muscle cross-sections at day 17. Note presence of RFP positive cells only in chimeras. Scale bar, 50µm. **(G)** Immunofluorescence staining for PAX7 in muscle cross-sections of a chimera at day 17, that has been subjected to host satellite cell ablation. White arrowheads point to PAX7^+^/RFP^-^ satellite cells. Scale bar 50µm. **(H)** A schematic illustrating strategy to assess if the RFP reporter is silenced in PAX7 expressing satellite cells. **(I)** Bright-field and fluorescence images of ITGA7^+^ satellite cell-derived myoblasts isolated from chimera muscles subjected to host satellite cell ablation. Scale bar, 100µm. **(J)** PCR for RFP in the indicated myoblast lines and conditions.

To assess which hypothesis is correct, we Fluorescence Activated Cell Sorting (FACS)-purified satellite cells using the established surface markers(44) CD45^-^/CD31^-^/SCA1^-^/ITGA7^+^ from muscles of chimeras subjected to host satellite cell ablation or non-injected controls (Figure S1C). Surprisingly, we detected both RFP positive and negative satellite cell populations and generated either RFP^+^ or RFP^-^ myoblast lines from chimeric muscles (Figures 1H and 1I). Importantly, PCR analysis for the RFP allele revealed that both the positive and negative RFP cell populations contained the RFP transgene, indicating that lentiviral vector silencing occurred in mESC-derived satellite cells (Figure 1J). Collectively, in this first trial we established a system to ablate host satellite cells in intraspecies chimeras and successfully produced satellite cells and myoblasts from donor-derived mESCs. However, lentiviral transgene silencing occurred in mESC-derived satellite cells, raising a need to find a genetic system that will allow to distinguish between host and donor-derived satellite cells.

### Exclusive production of gene-edited DMD iPSC-derived functional satellite cells in intraspecies chimeras

Given the encouraging results involving production of mESC-derived satellite cells in intraspecies chimeras, we next wished to test whether a similar approach may enable exclusive production of therapeutically-competent and gene-edited satellite cells from the well-established *Dmd^mdx^* mouse model(45). Specifically, we set out to explore whether we can produce and genetically correct *Dmd^mdx^*-iPSCs that carry a *Pax7-nuclear(n)GFP* satellite cell-specific genetic reporter, and then utilize *Dmd^mdx^; Pax7-nGFP* iPSCs to exclusively generate functional satellite cells in intraspecies chimeras following host satellite cell ablation (Figure 2A)(46).

**Fig. 2.**
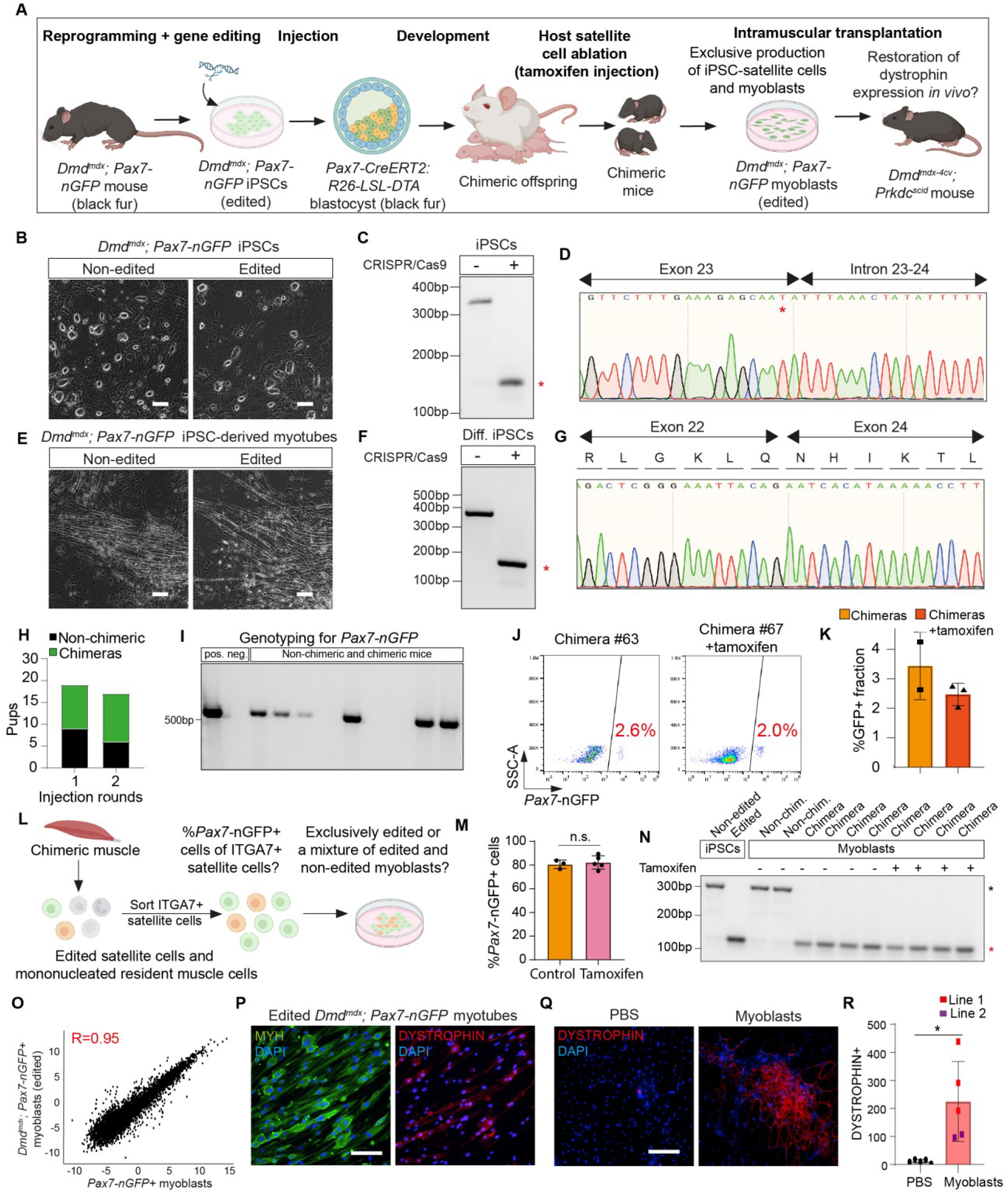
Functional muscle stem cells derived solely from edited *Dmd^mdx^; Pax7-nGFP* iPSCs in chimeras. **(A)** A schematic overview of experimental plan. **(B)** Representative bright-field images on the indicated cells. Scale bar, 500µm. **(C)** PCR products for *Dystrophin* amplified from DNA of non-edited (-) and edited (+) *Dmd^mdx^; Pax7-nGFP* iPSCs. Non-edited DNA formed a PCR fragment of 340bp, whereas gene editing leads to a shorter PCR product of 146bp (red asterisk). **(D)** DNA sequencing of the edited *Dystrophin* PCR product lacking a splice site. Red asterisk indicates the *mdx* mutation. **(E)** Myogenic differentiation of non-edited and edited *Dmd^mdx^; Pax7-nGFP* iPSCs into myotubes. Scale bar, 500µm. **(F)** PCR on cDNA of non-edited (396bp) and edited (183bp) *Dmd^mdx^; Pax7-nGFP* iPSC-derived myogenic cultures. The edited dystrophin band is marked by a red asterisk. **(G)** DNA sequencing of an edited dystrophin band, revealing successful exon skipping and reframing. **(H)** A graph showing chimera numbers. **(I)** Representative DNA genotyping for the *Pax7-nGFP* allele in in non-chimeric and chimeric offspring. **(J)** Flow cytometry analysis of Pax7-nGFP in the indicated animals and conditions. **(K)** Quantification showing the percentage of *Pax7-*nGFP^+^ cells in muscles derived from chimeras upon satellite ablation or control. N=2-3. **(L)** Strategy to investigate the proportion of *Pax7-*nGFP^+^ cells within the ITGA7^+^ satellite cell population. **(M)** Quantification of the percentage of *Pax7-nGFP*^+^ cells within the ITGA7^+^ satellite cell population derived from chimeras with or without tamoxifen treatment. N=3 for control, N=5 for the tamoxifen treated group. Statistical analysis was performed with percentage of GFP cells using ordinary one-way ANOVA, n.s., not significant. **(N)** PCR for dystrophin in ITGA7^+^-derived myoblasts from the indicated animals and conditions. Note that all myoblasts show only an edited band. **(O)** Scatterplot based on log2-normalized gene counts from bulk RNA-seq of the indicated samples. N=3 cell lines per group, p<2.2e-16. **(P)** Immunofluorescence staining for the indicated cells. Scale bar, 100µm. **(Q)** Immunofluorescence staining of cross-muscle sections *of tibialis anterior* (TA) muscles of the indicated conditions. Scale bar, 100µm. **(R)** Quantification of transplantation trials. Statistical analysis using paired t-test, N=5, ∗p ≤ 0.05, each dot represents an individual muscle.

As the first step, we crossed homozygous *Dmd^mdx^* female mice with homozygous *Pax7-nGFP* males and generated mouse embryonic fibroblast (MEF) lines. As the dystrophin gene is located on the X chromosome, all male MEF lines inherit the *Dmd^mdx^* mutation and are heterozygous for the *Pax7-nGFP* allele. Reprogramming to pluripotency was performed using a polycistronic *STEMCCA* cassette together with small molecule treatment (Figure S2A)(47, 48). Following manual picking and propagation of clones, we were able to establish *Dmd^mdx^; Pax7-nGFP* iPSCs that expressed well-known pluripotency markers (Figures S2B-E).

Next, we set out to correct the dystrophin mutation in exon 23 of *Dmd^mdx^; Pax7-nGFP* iPSC clones and employed a previously described CRISPR/Cas9 exon-skipping-based strategy that results in a restored reading frame (Figures S2F and S2G)(49). To this end, we utilized a previously reported single plasmid which encodes for Cas9, guide RNAs and a puromycin selection cassette (Figures S2F and S2G)(23). Transfection and antibiotic selection led to the generation of 24 DMD-iPSC sub-clones, of which two were correctly edited, and appeared indistinguishable from parental iPSCs (Figure 2B). We confirmed successful editing in these sub-clones at the DNA level by PCR and Sanger sequencing (Figures 2C and 2D). To unequivocally validate whether *Dmd^mdx^; Pax7-nGFP* iPSC sub-clones were successfully edited, we employed an established directed differentiation protocol of PSCs into muscle fibers *in vitro*(19, 50). This effort led to the generation of contractile muscle fibers from edited *Dmd^mdx^; Pax7-nGFP* iPSC sub-clones, which showed successful reframing of dystrophin at the mRNA level (Figures 2E-G). Furthermore, immunostaining revealed dystrophin^+^ myofibers solely in ESCs and edited *Dmd^mdx^; Pax7-nGFP* iPSCs subjected to the differentiation protocol, however not in unedited *Dmd^mdx^; Pax7-nGFP* iPSC-derived myogenic cultures subjected to this protocol (Figure S2H).

Based on these results, we proceeded to inject karyotypically normal and gene-edited *Dmd^mdx^; Pax7-nGFP* iPSCs into *Pax7-CreERT2: R26-LSL-DTA* blastocysts and produced 36 pups (Figures 2H and S2I). As both iPSCs and host blastocysts carry genes which encode for black fur coat color, we employed genotyping for the *Pax7-nGFP* allele to assess for chimerism, revealing this way that 21 out of 36 (58%) of the offspring were chimeric (Figures 2H and 2I). We then injected chimeras with tamoxifen between days 3-5 postnatally and harvested skeletal muscles from injected and non-injected chimeras at 5 weeks of age, aiming to assess the number of *Pax7*-nGFP*^+^* satellite cells with and without host satellite cell ablation (Figure 2A). Remarkably, we detected presence of *Pax7*-nGFP*^+^* satellite cells in chimeras following host satellite cell ablation (Figures 2J and S2J). Unexpectedly, we also detected approximately the same number of *Pax7*-nGFP^+^ satellite cells in non-injected chimeras, suggesting that cell ablation is not critical for substantial production of donor-derived satellite cells in chimeras (Figures 2J, 2K). FACS-purified satellite cells were then extracted from both injected and non-injected chimeras, giving rise to *Pax7*-nGFP^+^ myoblast lines (Figures S2J and S2K). Importantly, we confirmed that all of the *Pax7*-nGFP^+^ myoblast lines solely carried a correctly edited dystrophin gene (Figure S2L).

The observation that a comparable number of edited *Dmd^mdx^; Pax7-nGFP* satellite cells were generated in injected and non-injected chimeras promoted us to explore to what extent PAX7^+^ cell ablation may promote enhanced iPSC contribution to the satellite cell niche. To this end, we analyzed additional chimeras that have been treated with and without tamoxifen injections and FACS-purified satellite cells from their skeletal muscles using established satellite cell surface markers (CD45^-^/CD31^-^/SCA1^-^/ITGA7^+^)(44). We determined this way that most ITGA7^+^ satellite cells were GFP positive with and without tamoxifen administration (Figures 2L, 2M and S2M). We further plated CD45^-^/CD31^-^/SCA1^-^/ITGA7^+^ satellite cells and observed that nearly all myoblasts were GFP positive (Figures S2M and S2N). Most importantly, all ITGA7^+^ satellite cell-derived myoblast lines contained only the gene-edited dystrophin allele, suggesting that indeed all satellite cells were derived from edited *Dmd^mdx^; Pax7-nGFP* iPSCs (Figure 2N).

Next, we embarked on molecular and functional characterization of edited *Dmd^mdx^; Pax7-nGFP* myoblasts, confirming strong GFP expression in multiple myoblast lines (Figures S3A and S3B). Bulk RNA-seq analysis of FACS-purified edited *Dmd^mdx^*; *Pax7*-nGFP^+^ myoblasts revealed high expression of myoblast-related myogenic markers, and these cells were highly similar to FACS-purified PAX7^+^ myoblasts harvested from *Pax7-nGFP* mice(51) (Figures 2O and S3C). We then differentiated *Dmd^mdx^; Pax7-nGFP* myoblasts into myotubes by serum withdrawal and observed formation of elongated, multinucleated fibers that downregulated the *Pax7-nGFP* reporter (Figure S3D). PCR and cDNA sequencing analyses of differentiated myotubes revealed faithful correction of the mutation in dystrophin (Figures S3E and S3F). Notably, we detected dystrophin^+^ myotubes only in WT and edited *Dmd^mdx^; Pax7-nGFP* myotubes but not in unedited control (Figures 2P and S3G).

Finally, we explored whether edited *Dmd^mdx^; Pax7-nGFP* myoblasts can efficiently restore dystrophin expression *in vivo* in dystrophic muscles of immunodeficient *Prkdc^scid^; Dmd^mdx-4Cv^* mice, which were chosen due to the low rate of naturally-occurring revertant myofibers(52, 53). We transplanted 1 million edited *Dmd^mdx^; Pax7-nGFP* myoblasts into *tibialis anterior* (TA) muscles that have been pre-injured with cardiotoxin (CTX) injection to facilitate myoblast engraftment (Figures S3H and S3I). At 4 weeks post cell transplantation, we harvested the TA muscles and analyzed cross-muscle sections for presence of dystrophin expression. We observed a significant increase in dystrophin^+^ myofibers in dystrophic muscles transplanted with edited *Dmd^mdx^; Pax7-nGFP* myoblasts compared to PBS-injected controls (Figures 2Q and 2R). Altogether, we conclude that genetically-corrected *Dmd^mdx^; Pax7-nGFP* iPSCs can exclusively give rise to *bona fide* satellite cells in intraspecies chimeras, even without host satellite cell ablation. Myoblasts derived from these satellite cells can efficiently contribute to dystrophin restoration *in vivo*.

### Functional mouse satellite cells produced in interspecies rat-mouse chimeras

The capacity to generate genetically corrected satellite cells in intraspecies chimeras even without host satellite cell ablation prompted us to assess whether mouse satellite cells can be generated in another animal host. To address this objective, we chose rats as recipients, as xenogeneic cells and organs were previously produced in rat-mouse chimeras(37). We chose to inject edited *Dmd^mdx^; Pax7-nGFP* iPSCs into Sprague-Dawley (SD) rat blastocysts and assess production of satellite cells in rat-mouse adult chimeras (Figure 3A). We injected 8-12 edited *Dmd^mdx^; Pax7-nGFP* iPSCs into SD rat blastocysts, and embryos were transferred into the oviducts of foster rats. Collectively, this trial resulted in production of 4 rat-mouse chimeras out of 7 born pups (57.1%), as judged by patches of black fur. The extent of chimerism was varied, ranging between small fur patches to prominent contribution to fur black color (Figures 3B and S4A). Of note, most of these chimeras appeared healthy although one chimera, which showed the highest chimerism, was smaller in size and demonstrated body asymmetry and malocclusion (Figure S4A, top), in line with previous reports that documented abnormalities in interspecies chimeras with extensive xenogeneic contribution(54-56). We then opted to isolate and digest skeletal muscles from rat-mouse chimeras #2 and #4 and confirmed presence of the rat dystrophin allele and the mouse gene-edited dystrophin (Figures 3B, 3C and S4B). Flow cytometry analysis revealed presence of *Pax7*-nGFP^+^ satellite cells in the chimeras’ muscles, however less than in skeletal muscles harvested from young *Pax7-nGFP* mice (Figure 3D). Collectively, out of the four rat-mouse chimeras, we detected presence of *Pax7*-nGFP^+^ satellite cells in 3 chimeras (75%) (Figures 3D and S4C). We next decided to produce and characterize myoblast lines from rat-mouse chimeras #2 and #4, which showed robust *Pax7-nGFP* reporter expression *in vitro* (Figures 3D and 3E). DNA analysis confirmed the presence of only the corrected *dystrophin* gene and lack of rat DNA, as well as presence of the *Pax7-nGFP* transgene (Figures 3F, S4D and S4E). Additionally, myoblast mRNA showed reframing of dystrophin (Figures 3G and 3H). Of note, differentiation of myoblasts downregulated the *Pax7-nGFP* reporter and gave rise to dystrophin^+^ myotubes, unlike unedited control (Figures 3I and S4F).

**Fig. 3.**
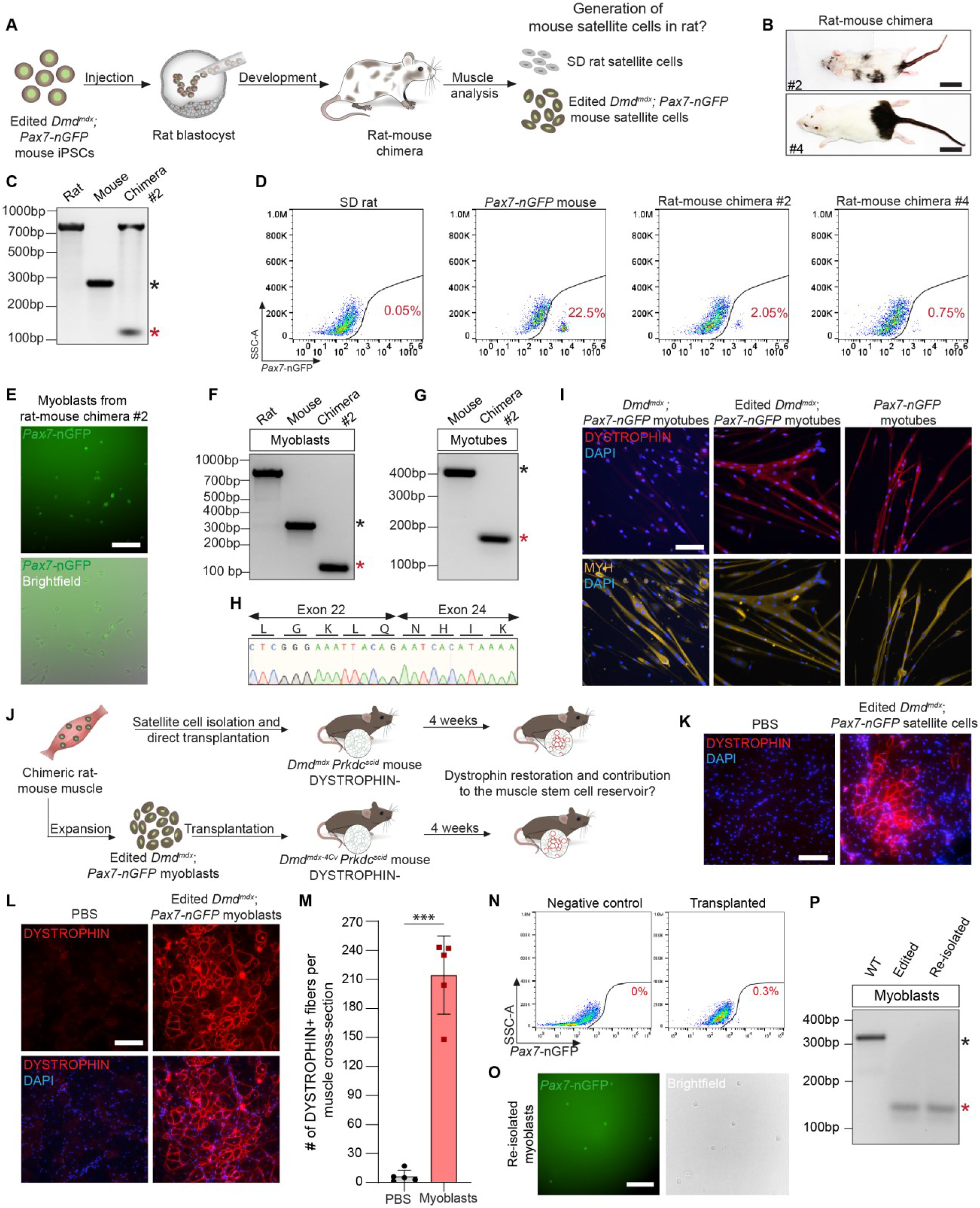
Production of edited *Dmd^mdx^; Pax7-nGFP* iPSC-derived functional mouse satellite in rat-mouse chimeras. **(A)** A schematic overview of experimental design. **(B)** Photos of rat-mouse chimeras at 7 weeks of age. Scale bar, 3.5cm. **(C)** PCR for rat and mouse dystrophin from digested muscle of the indicated animals. Black and red asterisks denote unedited (340bp) and edited (146bp) murine dystrophin, respectively. **(D)** Flow cytometry analysis of *Pax7-nGFP* expression in digested muscle of the indicated animal strains. **(E)** Representative images of *Dmd^mdx^; Pax7-nGFP* mouse myoblasts isolated from rat-mouse chimera #2 at P0. Scale bar, 100µm. **(F)** PCR for rat and mouse dystrophin of myoblasts isolated from the indicated animals. Black and red asterisks denote unedited and edited murine dystrophin, respectively. **(G)** PCR for mouse *Dystrophin* using cDNA of myotubes derived from edited *Dmd^mdx^; Pax7-nGFP* myoblasts produced in rat-mouse chimera vs. control. **(H)** Sanger sequencing reveals reframing of the dystrophin gene at the cDNA level of the myotubes analyzed in (G). **(I)** Immunofluorescence for dystrophin in edited *Dmd^mdx^; Pax7-nGFP* myotubes differentiated from rat-mouse chimera myoblasts vs. controls. Scale bar, 100µm. **(J)** A schematic overview of transplantation experiments. **(K)** TA muscle cross-section of *Dmd^mdx^; Prkdc^scid^* mice stained for dystrophin at 4 weeks after transplantation with satellite cells. Scale bar, 100µm. **(L)** *Dmd^mdx-4Cv^; Prkdc^scid^* TA muscle cross-sections stained for dystrophin at 4 weeks after transplantation with myoblasts. Scale bar, 100µm. **(M)** Quantification of dystrophin^+^ fibers. Statistical analysis was performed using paired t-test, N=5, ∗∗∗p ≤ 0.001, each dot represents an individual muscle. **(N)** Flow cytometry analysis of *Pax7-nGFP* expression in six TA muscles engrafted with myoblasts at 4 weeks after cell transplantation. **(O)** Representative images of myoblasts isolated in (N). Scalebar, 100µm. **(P)** PCR for mouse *Dystrophin* in myoblasts re-isolated from transplanted TA muscles shown in (N) and (O).

Last, we set out to explore whether xenogeneic satellite cells or derivative myoblasts produced in rat-mouse chimeras can restore dystrophin expression in DMD mice following intramuscular transplantation (Figure 3J). To this end, we FACS-purified *Pax7*-nGFP^+^ satellite cells from rat-mouse chimera #2 and injected about 10’000 satellite cells into CTX pre-injured TA muscles of *Prkdc^scid^; Dmd^mdx^* mice. Alternatively, at a later timepoint, we injected 1 million myoblasts at passage 7 that have been isolated from rat-mouse chimera #2 into CTX pre-injured TA muscles of *Prkdc^scid^; Dmd^mdx-4Cv^* mice (Figures 3J and S4G). At 1-month post transplantation, TA muscles were harvested, sectioned, and analyzed. Remarkably, we could detect areas of dystrophin^+^ myofibers in dystrophic muscles subjected to satellite or myoblast cell transplantation (Figures 3K and 3L). Quantification of these muscle cross-sections in comparison to a PBS-injected control revealed around 40 times more dystrophin^+^ myofibers following myoblast engraftment (Figure 3M and S4H). Surprisingly, we were also able to re-isolate transplanted myoblasts from recipient TA muscles by FACS-purification using the *Pax7-nGFP* reporter (Figures 3N and 3O). As further confirmation, a PCR analysis for the *dystrophin* gene on genomic DNA showed presence of only the edited band in these re-isolated myoblasts (Figure 3P). Together, these results demonstrate that mouse *Dmd^mdx^; Pax7-nGFP* iPSC-derived satellite cells or myoblasts produced in rat-mouse chimeras can efficiently restore dystrophin expression in muscles of DMD mice, and a fraction of cells can even remain as progenitors in the muscles enabling re-isolation of cells.

## Discussion

In this study, we report on production of genetically corrected and functional PSC-derived satellite cells in either mouse-mouse or rat-mouse chimeras. For intraspecies chimeras, we utilized an inducible ablation system of host *Pax7*^+^ cells to preferentially obtain ESC- or gene-edited DMD iPSC-derived satellite cells and derivative myoblasts capable of restoring dystrophin expression *in vivo* (Figure 4). Surprisingly, we observed efficient production of satellite cells even without an ablation system, which prompted us to investigate production of gene-edited iPSC-derived mouse satellite cells in rat-mouse chimeras. Strikingly, multiple rat-mouse chimeras contained appreciable numbers of donor-derived and gene-edited mouse satellite cells and derivative myoblasts that could efficiently restore dystrophin expression in DMD mice (Figure 4).

**Fig. 4.**
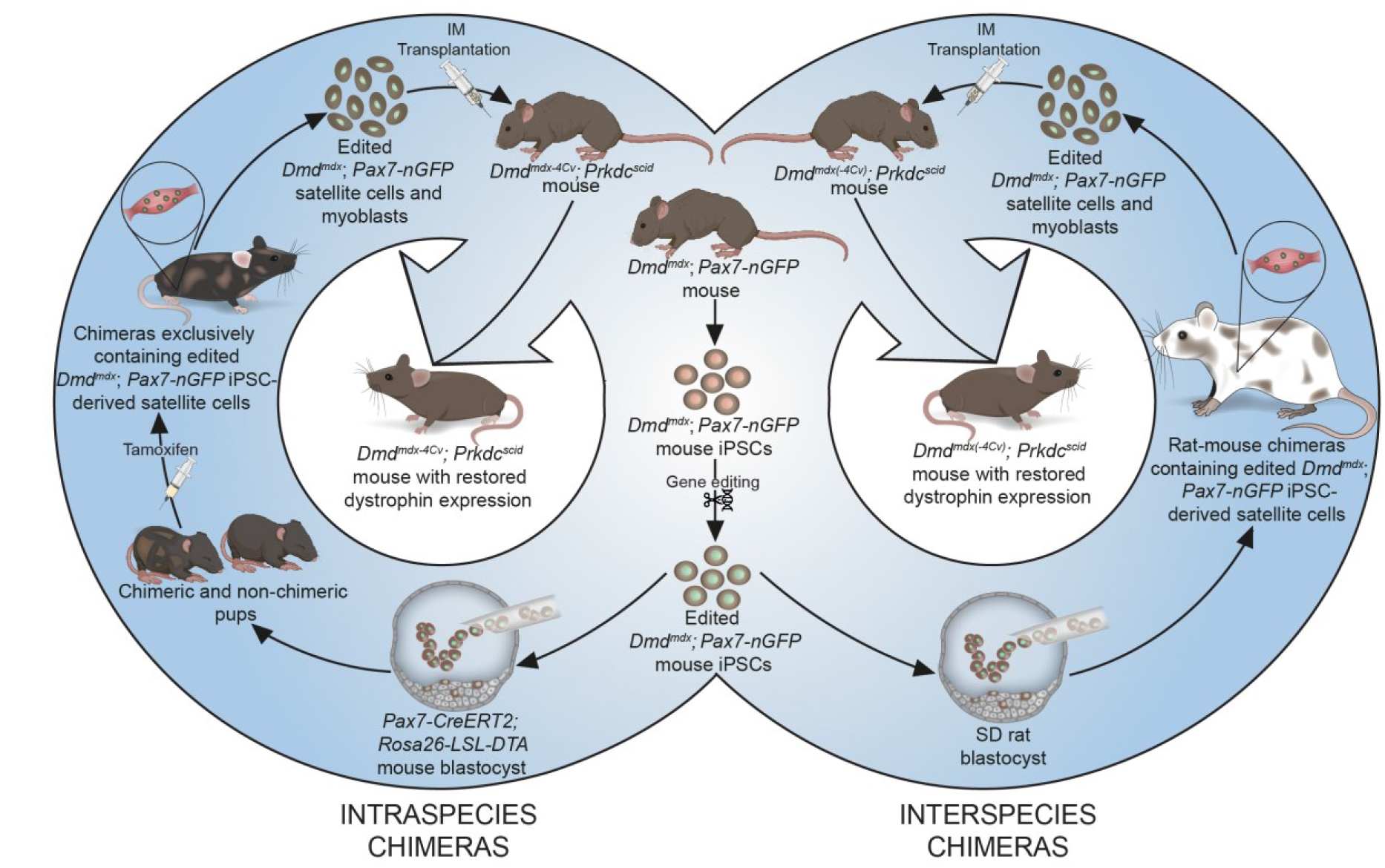
A schematic summarizing the study findings.

Our work raises the possibility that a similar approach may enable production of xenogeneic lineage-specific human adult stem cells in interspecies chimeras. In recent years, several papers reported on contribution of human PSCs to chimerism in mouse, pig and monkey embryos(57-61). However, adapting such a technique for production of human cells in animal chimeras is associated with ethical considerations and barriers, and most notably will require means to exclude production of undesired human cell types such as brain cells or gametes in human-animal chimeras(62-64). To this end, use of PSCs that carry a genetic mutation that precludes their differentiation into these cell types may provide a plausible solution as recently shown in mice(63).

Utilizing blastocyst complementation, a recent study reported on pig and human skeletal muscle formation by injection of pig PSCs or P53-null human iPSCs into pig embryos carrying triple knockout in *MYOD*, *MYF5* and *MYF6*, thus enabling PSC-colonization of the skeletal muscle lineage in chimeric embryos(60). However, assessing the contribution of human PSCs to the muscle stem cell compartment in born chimeras, or *in vitro* production of human myogenic cell lines from the interspecies embryos has not been documented(60). Moreover, one notable caveat for production of xenogeneic skeletal muscle tissue or organs in interspecies chimeras is the presence of animal host-derived endothelium, mesenchyme or other cell types, which may evoke an immunological response(34, 62). The approach reported in our study circumvents this major limitation as autologous muscle stem cells can be FACS-purified in considerable numbers from interspecies chimera muscles and transplanted into patients, without lingering animal cells. Another advantage for the approach detailed in our study is that the PSCs were differentiated *in vivo*, thus mitigating any potential future risk for residual PSCs to form teratomas in patients as recently reported for *in vitro* PSC-derived cells(65). Last, as iPSCs differentiate into satellite cells in postnatal chimeras, this approach ensures the generation of adult muscle stem cells, in comparison to myogenic cells generated from PSCs via directed differentiation *in vitro*, which may carry embryonic attributes(66).

To close, our study demonstrates a proof-of-principle for combining cellular reprogramming, genome engineering and *in vivo* PSC differentiation to produce therapeutically-competent muscle stem cells in a xenogeneic chimera host. In respect to implications for human therapy, further work is certainly warranted to address the many hurdles associated with generating human cells in animals. However, in the event such roadblocks are overcome, we envision this work may pave the way for producing autologous human satellite cells in large animals for the treatment of muscle diseases.

## Materials and Methods

### Animals

Mice and rats used in this study were housed in Allentown cages, under standard conditions at room temperature 23°C and a relative humidity of 50-60%, with a 12-h light-dark cycle. All animals had *ad libitum* access to food and water. The following mouse strains from Jackson Laboratories were used: C57BL/10ScSn-*Dmd^mdx^*/J (Stock No: 001801); B10ScSn.Cg-*Prkdc^scid^ Dmd^mdx^*/J (Stock No: 018018); B6; 129S-*Gt(ROSA)^26Sortm1.1Ksvo^*/J (Stock No: 023139); B6.Cg-*Pax7^tm1(cre/ERT2)Gaka^*/J (Stock No: 017763). Additionally, the previously reported strains were used: *Tg:Pax7-nGFP/C57BL6;DBA2* mice (a kind gift from Dr. Shahragim Tajbakhsh)(46) for production of MEFs for iPSC reprogramming and NOD.Cg-Prkdc^scid^l2rgtm1Wjl/SzJ (005557)x B6Ros.Cg-Dmd^mdx-4Cv^/J (002378) as recipient mice for intramuscular myoblast transplantations. Swiss webster mice and Sprague-Dawley rats were purchased from Janvier, France. The present study was approved by the Federal Food Safety and Veterinary Office, Cantonal veterinary office (Zurich) and granted animal experiment license numbers ZH246/18, ZH177/18 and FormG-135.

### Cell culture

Mouse embryonic fibroblasts (MEFs) were isolated from E13.5 mouse embryos and grown in ‘MEF medium’ containing DMEM (41966029, Thermo Fisher Scientific) supplemented with 10% FBS (10270106, Thermo Fisher Scientific), 1% MEM Non-Essential Amino Acids Solution (100X) (11140050, Thermo Fisher Scientific), 1% Penicillin-Streptomycin (10,000 U/mL) (15140122, Thermo Fisher Scientific) and 0.1% Gibco 2-Mercaptoethanol (21985023, Thermo Fisher Scientific). HEK-293T cells were grown in ‘MEF medium’. MEF reprogramming and iPSCs culture was performed in MES medium consisting of DMEM (41966029, Thermo Fisher Scientific), 1% GlutaMAX Supplement (35050061, Thermo Fisher Scientific), 1% Penicillin-Streptomycin (10,000 U/mL) (15140122, Thermo Fisher Scientific), 1% MEM Non-Essential Amino Acids Solution (100X) (11140050, Thermo Fisher Scientific), 0.1% Gibco 2-Mercaptoethanol (21985023, Thermo Fisher Scientific), 15 % FBS (10270106, Thermo Fisher Scientific) supplemented with 1000U/ml mLIF (#PG-A1140-0010, PolyGene Transgenetics). iPSCs and ESC were cultured in ‘Enhanced medium’ consisting of MES medium supplemented with 0.5µmol/l VPA (P4543-10G, Sigma-Aldrich), 1.5µmol/l CGP77675 (SML0314, Merck) and 3µmol/l CHIR99021 (4423, Tocris Bioscience) for 5 days prior to blastocyst injections. ‘Myoblast medium’ for satellite cells and myoblast consisted of 50% DMEM (41966029, Thermo Fisher Scientific), 50% F-10 medium (22390025, Thermo Fisher Scientific), 10% Horse serum (16050122, Thermo Fisher Scientific), 20% FBS (10270106, Thermo Fisher Scientific), 1% Penicillin-Streptomycin (10,000 U/mL) (15140122, Thermo Fisher Scientific) and 10ng/ml basic FGF (233-FB-500, R&D Systems).

Myoblast were differentiated in ‘Differentiation medium’ containing DMEM (41966029, Thermo Fisher Scientific) supplemented with 2% Horse serum (16050122, Thermo Fisher Scientific) and 1% Penicillin-Streptomycin (10,000 U/mL) (15140122, Thermo Fisher Scientific). All cells were passaged using Gibco Trypsin-EDTA (0.05%) (25300054, Thermo Fisher Scientific) and tested for mycoplasma (LT07-318, Lonza). All cells were maintained at 37°C in a 5% CO_2_ incubator.

### Generation of iPSCs

To generate iPSCs, male MEFs were transduced with lentiviral vectors FUW-M2rtTA (Addgene plasmid #20342) and LV-mSTEMCCA-1(48) combined with 5ug/ml Polybrene (TR-1003-G, Sigma-Aldrich) on two consecutive days in MEF medium without Penicillin-Streptomycin. 20’000 transduced MEFs were then seeded into one well of a 6 well plate in ‘MES medium’ supplemented with 2ug/ml Doxycycline (D9891-5G, Sigma-Aldrich), 3μM CHIR99021 (4423/50, R&D Systems) and Ascorbic acid (A4403-100MG, Sigma-Aldrich) at the final concentration of 50 µg/ml to initiate reprogramming. Reprogramming medium was changed daily until iPSC colonies appeared. Dox-independent iPSCs colonies were picked and transferred onto γ-irradiated CF-1 fibroblasts and expanded in ‘MES medium’.

### Lentivirus preparation

Frozen lentiviral stocks were generated by transfecting 70% confluent HEK-293T cells in 15cm cell culture dishes in ‘MEF medium’ together with a solution consisting of 1ml 150mM NaCl (1.06404.1000, VWR), 1ml Polyethylenimine (23966-1, Polysciences), pLV delta 8.9 (16.5μg), pLV VSVG Envelope (11μg) and the transfer vector (22μg). 24 hours after transfection, the cell medium was changed to regular ‘MEF medium’. Supernatant was collected, filtered using a 0.45μm filter and stored at 4°C 48h and 72h after transfection. 0.25ml of cold PEG-it Virus Precipitation Solution (LV810A-1-SBI, System Biosciences) was added per 1ml of lentiviral-containing supernatant and kept cool at 4°C overnight. Next, the mixture was centrifuged at 1500g for 30min at 4°C, resuspended in PBS containing 25mM HEPES (15630056, Thermo Fisher Scientific) and stored at -80°C for future use.

### Gene editing

iPSCs were transfected as single cells using Lipofectamine CRISPRMAX Cas9 Transfection Reagent (CMAX00003, Thermo Fisher Scientific) according to the manufacturer’s instructions with pRP[CRISPR]-EGFP/Puro-hCas9-U6>(long left)-U6>(long right) (Vector builder, VB190118-1126uvv). 1 day after transfection, puromycin (A1113803, Thermo Fisher Scientific) was added at a final concentration of 1 µg/ml for 3-4 days for selection. After recovery in puromycin-free medium, single iPSC colonies were picked and expanded. 24 colonies were picked and characterized. 2 out of these showed the expected editing with an additional 6 bp intronic deletion that didn’t impact the final outcome.

### iPSC differentiation

iPSC differentiation towards myogenic cells was performed as described by Chal et al(19, 50).

### Karyotyping

KaryoMAX Colcemid Solution (15212012, Thermo Fisher Scientific) was added to the iPSCs at a final concentration of 100 ng/ml. After a 5-hour incubation at 37°C, cells were harvested, centrifuged, and supernatant was removed to 1 ml. Cells were re-suspended by vortexing at low setting and 5ml of pre-warmed hypotonic solution (0.56g KCl+0.5g sodium citrate in 200ml H2O) were added dropwise while mixing. The cell suspension was then incubated at 37°C and after 30min, 2.5ml of fixative (Methanol:Acetic acid = 3:1) were added. The mixture was centrifuged, supernatant removed, and the cell pellet was re-suspended in 2ml of fixative while vortexing and incubated at room temperature for 5 min. This step was repeated 3 times. Next, the cell pellet was re-suspended in 1ml fixative and 400µl of this cell suspension was dripped on a polarized microscope slide from a height of 150cm. After the slides dried, they were immersed in Giemsa solution for 7 min, put into Gurr’s buffer (0.469 g NaH_2_PO_4_ + 0.937 g Na_2_HPO_4_ in 1 l of H_2_O) for 2 min and then washed with water. Slides were let to dry and examined. For analysis, 30 cell spreads were randomly selected and chromosomes were counted.

### Blastocyst injection

Blastocyst injections and embryo transfer were performed in house, abiding to all legal rules of the Federal Food Safety and Veterinary Office, Cantonal veterinary office (Zurich) and an animal experimental license (FormG-135 and ZH246/18). On the injection day, cells were pre-plated and then kept on ice in until injection. For mouse intraspecies blastocyst injections, mice aged 3-4w were superovulated via intraperitoneal injection of 5IU PMSG (ProSpec). About 46-48h later, the mice were injected again with 5IU hCG (ProSpec), to induce ovulation, and were paired with stud males. The following morning the females were separated, euthanized after 48 hours, the morulae flushed from the oviduct using M2 Medium (M7167, Sigma-Aldrich) and incubated overnight at 37°C and 5% CO2. For mouse to rat interspecies blastocyst injections, SD rats (aged 8-15w) were synchronized via injection of 40µg of LHRHa (L4513, Sigma-Aldrich). About 94-96h later, the rats were paired with stud males. After 116-120h, the blastocysts were flushed in M2 medium. Injections were carried out in droplets of M2 medium covered with Mineral Oil (Sigma-Aldrich,

M8410-1L). Typically, 8-12 iPSCs or ESCs cells were injected per blastocyst. Successfully injected blastocysts were transferred to a 60ml center well organ culture dish containing CO2-buffered culture medium (EmbryoMax KSOM medium (MR-106-D, Sigma-Aldrich) for mouse blastocysts, rat KSOM (CSR-R-R148, Cosmo Bio) for rat blastocysts) and kept in an incubator at 37°C and 5% CO2. The injected mouse blastocysts were transferred to the uteri of 2.5dpc pseudo-pregnant female mice on the same day, or into uteri of 3.5pdc pseudo-pregnant female SD rats in case of rat blastocysts. Injections were performed with a Narishige MTK-1 hydraulic micromanipulator (Narishige), combined with a CellTram oil and a PiezoXpert (both Eppendorf). Presence of chimerism in intraspecies or interspecies chimeras was first assessed using visual examination of fur coat color. Further, chimerism was also assessed by genotyping for the RFP transgene or the *Pax7-nGFP* reporter. All the rat-mouse chimeras had a rat *Pax7*+/- background.

### Satellite cell ablation

A tamoxifen inducible ‘*Pax7-CreERT2: R26-LSL-DTA*’ system was used to ablate host satellite cells. Mice received an intraperitoneal injection of 50μl of 1mg/ml tamoxifen (T5648-1G, Sigma-Aldrich) in corn oil (C8267, Sigma-Aldrich) every day at P3-P5 followed by bi-weekly injections /75 mg tamoxifen/kg body weight).

### Intramuscular transplantation of myoblasts and satellite cells

Either *B10ScSn.Cg-Prkdc^scid^ Dmd^mdx^/J* mice or *NOD.Cg-Prkdc^scid^ /SzJ; B6Ros.Cg-Dmd^mdx-4C^v/J* received an injection of 50μl of 10μM Cardiotoxin (L8102, Latoxan) into the hindleg *tibialis anterior (TA)* muscles to enhance engraftment of the cells to be injected the day after. For myoblast transplantation, myoblasts were harvested, and 1’000’000 cells were resuspended in 20µl PBS. For satellite cell transplantation, 10’000 satellite cells were centrifuged at 650g for 5 min immediately after FACS and resuspended in 20µl PBS. This cell mixture was then injected craniocaudally into the pre-injured TA muscle (left leg) using an insulin syringe (324824, BD) and the cell suspension was slowly released upon retraction of the cannula. As a control, PBS was injected into the right hindleg pre-injured TA muscle. Mice were euthanized and TAs harvested 27-28 days after cell transplantation. Fifteen *NOD.Cg-Prkdc^scid^ /SzJ; B6Ros.Cg-Dmd^mdx-4C^v/J* recipient mice were used for the transplantation experiment using three different myoblast lines (two from intraspecies chimeras and one from interspecies chimera), but only ten of these mice showed engraftment. Two *B10ScSn.Cg-Prkdc^scid^ Dmd^mdx^/J* mice were used for direct transplantation of satellite cells, which engrafted in only one of them.

### Muscle embedding and processing

Harvested TAs were placed onto a cork covered with 10% Tragacanth (G1128, Sigma-Aldrich), and frozen for 30-60s in 2-Methylbutane (3927.1, Carl Roth) pre-cooled in LN_2_. The samples were added to liquid nitrogen for 1min and then stored at -80°C. Frozen muscles were cryo-sectioned into 10μm thick cross-sectional muscle sections and stored at -80°C.

### Genomic DNA isolation and PCR

Isolation of genomic DNA from cells was performed using the DNeasy Blood & Tissue Kit (69504, Qiagen) according to the manufacturer’s instruction. DNA for PCR from muscle lysates was obtained by direct lysis of muscle slurry generated by muscle mincing followed by 90 min incubation in digestion solution containing 2mg/ml Collagenase Type 2 (17101015, Thermo Fisher Scientific) in DMEM (41966029, Thermo Fisher Scientific) at 37°C using DirectPCR Lysis Reagent (mouse tail) (VIG102-T, Viagen Biotech). PCR for *Pax7-nGFP* was performed using primer Pax7nGFP.F/ Pax7nGFP.R and GoTaq G2 Hot Start Green Master Mix (M7423, Promega) via the following program: 94°C 2min, 30x (94°C 30s, 65°C 30s, 72°C 30s), 72°C 5min. Dystrophin gene editing was verified by PCR for mouse dystrophin using primers Dmd_i22-i23.F/ Dmd_i22-i23.Rand GoTaq G2 Hot Start Green Master Mix (M7423, Promega) via the following program: 94°C 5min, 30x (94°C 30s, 55°C 30s, 72°C 30s), 72°C 5min. PCR products were separated on a 1.5-2% agarose gel (7-01P02-R, BioConcept) dissolved in TAE buffer (3-07F03-I, BioConcept) and visualized using GelRed Nucleic Acid Stain (Cat# 41003, Biotium). Presence of rat cells in myoblast culture/muscle lysate was assessed by PCR for rat dystrophin using primers Rat Dmd i22-i23.F/ Rat Dmd i22-i23.R and GoTaq G2 Hot Start Green Master Mix (M7423, Promega) via the following program: 94°C 5min, 30x (94°C 30s, 57°C 1min, 72°C 30s), 72°C 5min.

### RNA Extraction and cDNA synthesis

RNA isolation was performed using Qiagen RNeasy kit, with 15min DNase digest (74104, Qiagen). The concentration was measured by a Tecan Spark 10M. cDNA was synthesized with the High-Capacity cDNA Reverse Transcription Kit (4368814, Thermo Fisher Scientific) using 1µg of RNA according to the manufacturer’s instruction.

### RT-PCR and RT-qPCR on cDNA

To verify *Dystrophin* gene editing on mRNA derived cDNA, RT-PCR was performed with primers Dmd_e22-e24.F / Dmd_e22-e24.R using GoTaq G2 Hot Start Green Master Mix (M7423, Promega) via the following program: 94°C 5min, 35x (94°C 30s, 55°C 30s, 72°C 30s), 72°C 5min. PCR products were run on a 1.5-2% agarose gel (7-01P02-R, BioConcept) in TAE buffer (3-07F03-I, BioConcept) and imaged using GelRed Nucleic Acid Stain (41003, Biotium). For RT-qPCR, the Applied Biosystems PowerUp SYBR Green Master Mix (A25741, Thermo Fisher Scientific) was used, with a final cDNA concentration of 5ng/μl according to the manufactures manual and primer sets mGapdh.F/mGapdh.R, mNanog.F/mNanog.R, mOct4.F/mOct4.R, mSox2.F/mSox2.R, Pkg.F/Pkg.R. RT-qPCR reactions were run on a 384-well block QuantSudio 5 System with comparative CT standard run mode and the following cycling conditions: Hold stage 50°C 2min-95°C 10min; PCR stage: 95°C 15s-60°C 1min, 40x; Melt Curve stage: 95°C 15s-60°C 1min-95° 15s. All reactions were done in triplicate and Ct values were obtained. Data was analyzed using 2-ΔΔCt method and mouse *Pgk* and *Gapdh* served as housekeeping control genes.

### Sequencing

Dystrophin DNA and cDNA PCR products were run on a 1.5-2% agarose gel (7-01P02-R, BioConcept) in TAE buffer (3-07F03-I, BioConcept) and products were extracted using the QIAquick Gel Extraction Kit (28706, Qiagen) and Sanger sequenced by Microsynth (Balgach, Switzerland) and evaluated with Benchling (retrieved from https://benchling.com in 2020) or SnapGene Viewer 3. The following primers were used for sequencing: Dmd_e22-e24.F (for sequencing of PCR products obtained from myoblast cDNA), Dmd_i22-i23.F (for sequencing of PCR products obtained from iPSC genomic DNA).

### Alkaline phosphatase test

Alkaline phosphatase test was performed using the Leukocyte Alkaline Phosphatase Kit (86R-1KT, Sigma-Aldrich). First, alkaline dye mixture was prepared as described in manufacturer’s protocol. Next, the fixing step was adopted for cells in a culture dish: 0.5ml of fixative solution prepared as described by manufacturer was added to each well of the 6-well plate with the cells. After 1min incubation at room temperature, cells were washed with PBS twice. Finally, 1ml of alkaline dye mixture per well of 6-well plate was added and plates were incubated for 15 min at room temperature in the dark. Alkaline dye mixture was aspirated, cells were washed twice with PBS, covered with 0.5 ml of PBS and imaged.

### Immunofluorescent staining

Cells cultured in 6 well plates were washed with PBS and fixed with 4% Paraformaldehyde (11400580, Fisher Scientific) in PBS for 5min at room temperature (RT). After two PBS washing steps, ‘blocking solution’ made up of PBS supplemented with 2% BSA (A1391, AppliChem) and 1% Triton X-100 (9002-93-1, Sigma-Aldrich) was added to the fixed cells for 30min at RT. Primary antibodies diluted in ‘blocking solution’ containing 0.2-1% Triton-X-100 were then added for 1 h at RT. Cells were washed twice with PBS and incubated with secondary antibodies and DAPI (62248, Thermo Fisher Scientific) in ‘blocking solution’ for 1h at RT. Cells were again washed twice with PBS and covered with ProLong Gold Antifade Mountant (P36934, Thermo Fisher Scientific) to prevent photobleaching before storage at 4°C.

TA muscle sections on microscope slides (J1800AMNZ, Epredia) were fixed with 4% Paraformaldehyde (11400580, Fisher Scientific) in PBS for 5min at RT and washed twice with PBS. ‘Blocking solution’ consisting of PBS supplemented with BSA (Cat. # A1391, AppliChem) and 0.2% Triton X-100 (Cat. #9002-93-1, Sigma-Aldrich) was added for 15min at RT. Primary antibodies diluted in ‘blocking solution’ were added for 1h at RT, after which the sections were washed twice with PBS and incubated for 30min with a secondary antibody and DAPI (62248, Thermo Fisher Scientific) diluted in ‘blocking solution. Sections were washed twice with PBS and covered with a few drops of ProLong Glass Antifade Mountant (P36980, Thermo Fisher Scientific) and a coverslip and stored at 4°C.

The following primary antibodies were used in this study: SOX2 Polyclonal Antibody (48-1400, Thermo Fisher Scientific), rat anti-mouse NANOG conjugated eFluor 660 (50576182, Thermo Fisher Scientific), OCT4 monoclonal antibody (9B7) (MA1-104, Thermo Fisher Scientific) and anti-Dystrophin antibody (Cat.#ab15277, Abcam), all diluted 1:200, Myosin Heavy Chain antibody (MAB4470, R&D Systems) diluted 1:500, anti-Laminin antibody (ab11575) (Cat.#ab11575, Abcam) and Human/Mouse/Rat/Chicken anti-PAX7 antibody (MAB1675, R&D Systems, 5μg/ml), both diluted 1:100. Secondary antibodies used were: goat anti-rabbit IgG 488 (A11008, Thermo Fisher Scientific), goat anti-mouse IgG1 488 488 (A21121, Thermo Fisher Scientific), goat anti-mouse IgG2b 647 (A21242, Thermo Fisher Scientific), goat anti-Mouse IgG1 647 (A-21240, Thermo Fisher Scientific), anti-rabbit IgG 647 (A31573, Thermo Fisher Scientific), anti-mouse 488 (A-21141, Thermo Fisher Scientific). All secondary antibodies were diluted 1:400.

### Satellite cell isolation and FACS

A Sony SH800S Cell Sorter was utilized for flow cytometry analysis and cell sorting for satellite cell isolation using Pax7-nGFP reporter, cell surface markers and for the purification of GFP positive myoblasts. For satellite cell isolation, skeletal muscles were harvested, minced and centrifuged in PBS at 350g for 3 min. The tissue pellet was then resuspended in digestion solution containing 2mg/ml Collagenase Type 2 (17101015, Thermo Fisher Scientific) in DMEM (41966029, Thermo Fisher Scientific) and incubated for 90min. This was followed by a 30min incubation step in a digestion solution consisting of 14ml F-10 (Thermo Fisher Scientific, 22390025) supplemented with 10% horse serum (Thermo Fisher Scientific, 16050122), 1ml of 0.2% Collagenase Type II (Thermo Fisher Scientific, 17101015) and 2.5ml of 0.4% Dispase II (Thermo Fisher Scientific, 17105041). Both incubation steps were performed in a shaking 37°C water bath. An 18 gauge needle syringe was then used to dislodge the cells from the fibers, followed by filtering the cells with a 100μm, 70μm and either 30μm or 40μm cell strainers. The cell pellet was resuspended in ‘FACS buffer’ consisting of PBS and 2% FBS (10270106, Thermo Fisher Scientific) and kept on ice until FACS. Satellite cells were FACS-purified using either Pax7-nGFP reporter or the following combinations of cell surface markers: APC anti-mouse Ly-6A/E (Sca-1) Antibody (108111, BioLegend), mouse Integrin alpha 7 Alexa Fluor 750-conjugated Antibody (FAB3518S, R&D Systems), APC anti-mouse CD45 Antibody (103111, BioLegend), APC anti-mouse CD31 Antibody (102409, BioLegend). In each FACS experiment, DAPI (62248, Thermo Fisher Scientific) was used to exclude dead cells. For GFP based sorting, GFP positive cells were sorted using the EGFP channel. For surface marker-based sorting, all SCA1^+^, CD45^+^ and/or CD31^+^ cells were excluded using APC channel. ITGA7^+^ were sorted from the remaining population and analyzed for GFP expression.

### RNA sequencing

RNA sequencing was performed at the FGCZ on an Illumina NovaSeq instrument and library was prepared according to Illumina Truseq mRNA protocol. The sequencing reads were analyzed using the SUSHI framework (67, 68) developed at the FGCZ. After the quality control (adapter and low-quality base trimming) with fastp v0.20(69), raw reads were pseudo-aligned against the reference mouse genome assembly GRCm39 and gene expression level (GENCODE release 26) was quantified using Kallisto v0.46.1(70). Genes were considered to be detected if they had at least 10 counts in 50% of the samples. Pax7-nGFP MEFs and Pax7-nGFP^+^ myoblasts samples from the GEO data set GSE169053 were re-processed in the same way (51). TPM (Transcripts per Million mapped reads) was used as the unit for normalized gene expression.

### Statistical analysis

Statistical analysis was performed with GraphPad Prism (Version 9.2.0, GraphPad Software) and presented as mean ± SD. Values of p<0.05 were considered statistically significant. Differences were evaluated using Student’s t-test (two sided) or one-way or two-way ANOVA.

### Study approval

The present study was approved by the Federal Food Safety and Veterinary Office, Cantonal veterinary office (Zurich) and granted animal experiment license numbers ZH246/18, ZH177/18 and FormG-135.

## Author contributions

Conceptualization: SD, AL, JZ, OBN

Experiments intraspecies part: SD, AL, NB

Experiments interspecies part: AL, JZ, NB, GB

Chimera production: MTS

RNA sequencing analysis: AG

Writing – original draft: SD, AL, JZ, OBN

## Acknowledgments

We are thankful to Inseon Kim and Xhem Qabrati for their feedback and fruitful discussions. We further wish to thank Pjeter Gjonlleshaj for his help with rat superovulation and pairing. We thank Dr. Konrad Hochedlinger for providing the M2rtTA and STEMCCA lentiviral cassettes and Dr. Shahragim Tajbakhsh for providing the *Pax7*-nGFP mouse strain. We are also grateful to the Functional Genomics Center Zurich (FGCZ) for their assistance with bulk RNA sequencing. A few of the graphical schematics were created with BioRender.com under a paid license, other schematics were generated by Veronique Juvin from SciArtWork.

## Tables

**Table 1:**
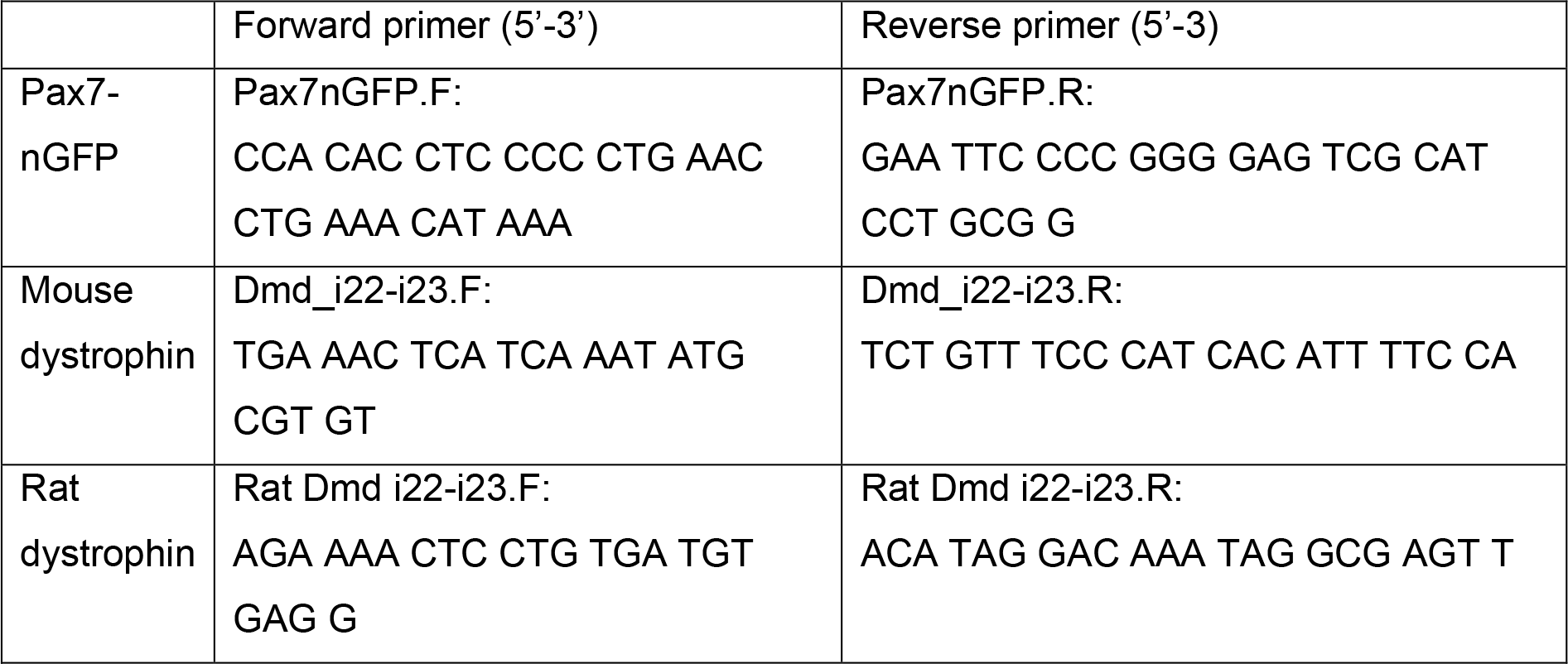
PCR primers

**Table 2:**
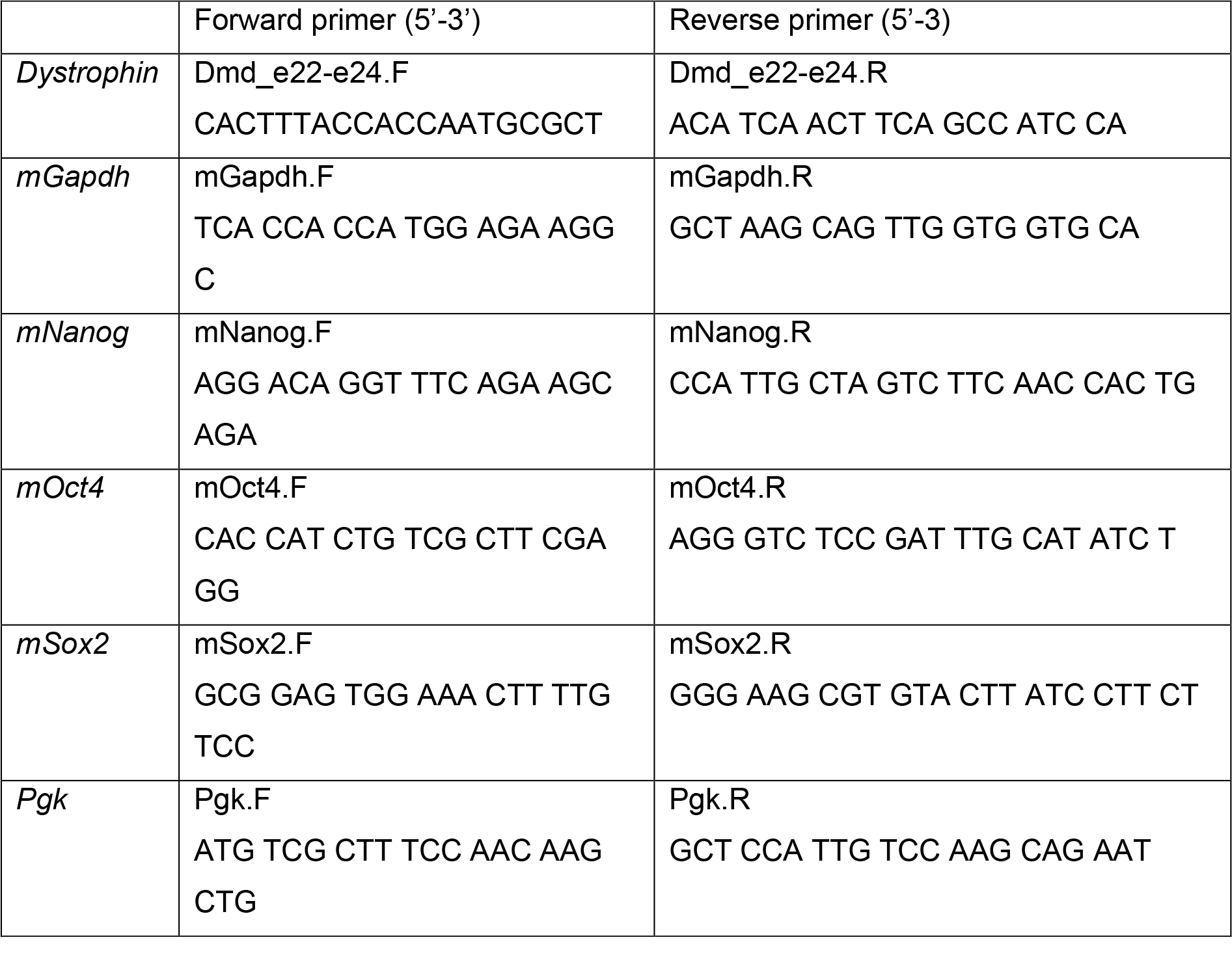
RT-qPCR primers

