## Supplementary figures for "Interspecies generation of functional muscle stem cells"

**Figure S1**

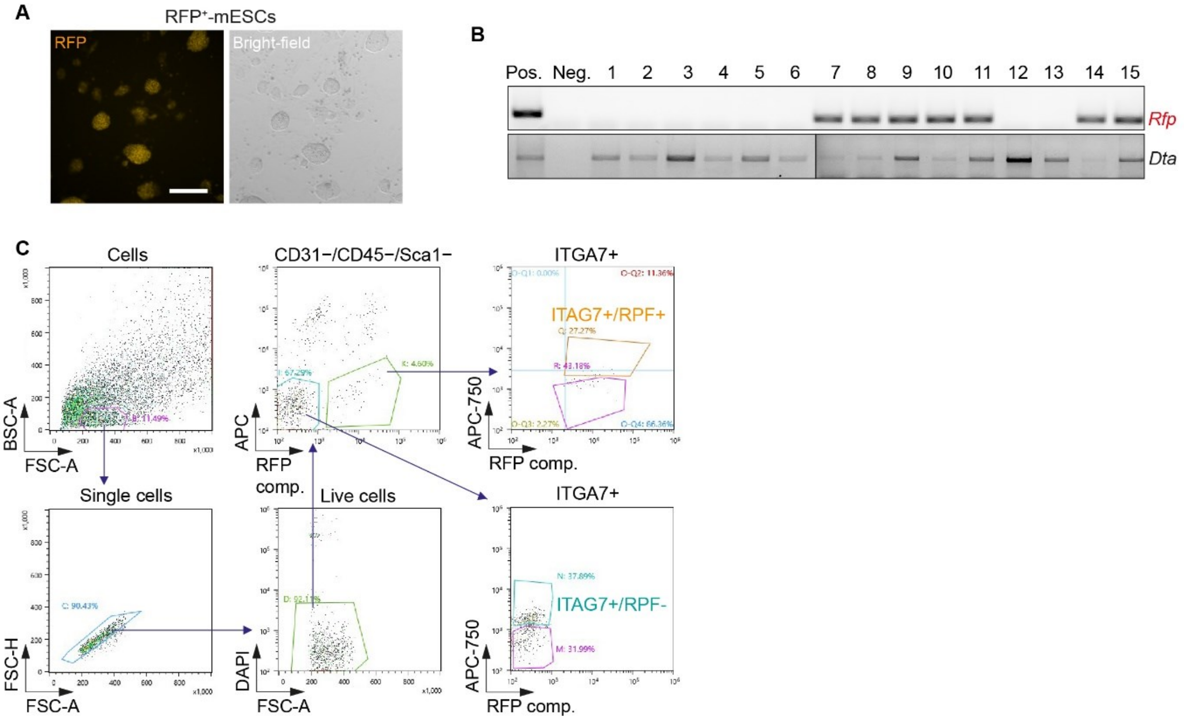


**Fig. S1.**

**(A)** Microscopy images of RFP$^+$-mESCs. Scalebar, 100µm. **(B)** PCR of DNA extracted from ear clips of non-chimeric and chimeric mice for the *Rfp* transgene and *Rosa26-LSL-DTA* allele. **(C)** FACS plots displaying satellite cell sorting strategy.

**Figure S2**

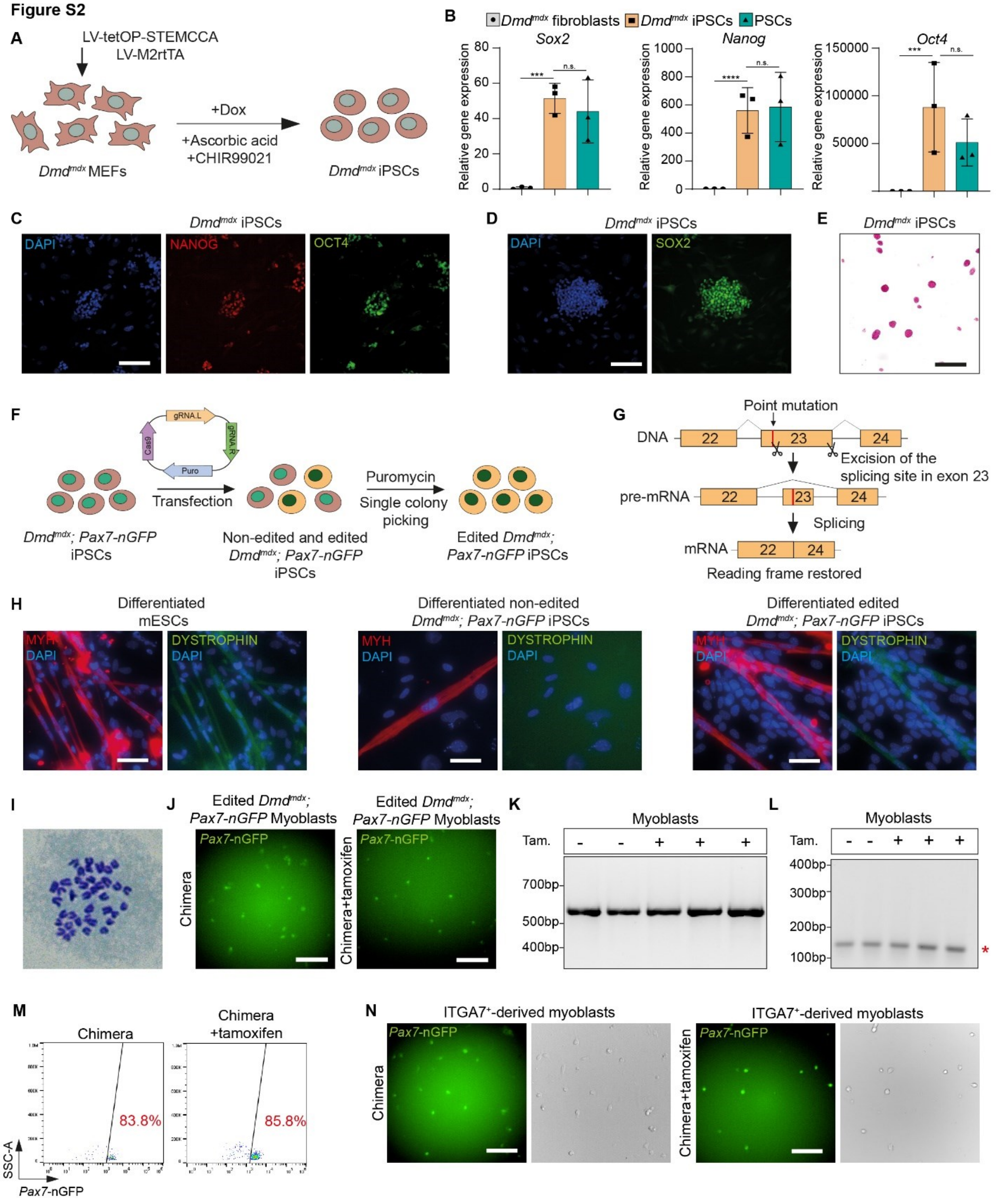


**Fig. S2.**

**(A)** Schematic overview. MEFs, mouse embryonic fibroblasts; dox, doxycycline. **(B)** Quantitative real-time PCR for the indicted genes. N=3 different lines, error bars denote SD. Statistical analysis was performed with delta Ct values using ordinary one-way ANOVA. $***p \leq 0.001$, $****p \leq 0.0001$, n.s., not significant. **(C)** Immunofluorescence images for the indicated proteins. Scalebar, 100µm. **(D)** *Dmd*$^{mdx}$ iPSCs showing expression of SOX2. Scalebar, 100µm. **(E)** Alkaline phosphatase staining in *Dmd*$^{mdx}$ iPSCs. Scalebar, 500µm. **(F)** Overview of gene editing of *Dmd*$^{mdx}$*; Pax7-nGFP* iPSCs. **(G)** Strategy showing CRISPR/Cas9 induced gene editing of exon 23 in the dystrophin gene. **(H)** Immunofluorescence staining for DYSTROPHIN in the indicated samples. Scalebar, 50µm. **(I)** Karyotype of edited *Dmd*$^{mdx}$*; Pax7-nGFP* iPSCs. **(J)** Edited *Dmd*$^{mdx}$*; Pax7-nGFP* myoblasts FACS-purified from muscle tissue of the indicated chimeras. Scalebar, 100µm. **(K)** PCR gel showing genotyping of the *Pax7-nGFP* allele in chimera-derived myoblasts that have been treated with or without tamoxifen (tam). **(L)** PCR for *Dystrophin* using DNA of edited *Dmd*$^{mdx}$*; Pax7-nGFP* myoblasts that have been FACS-purified from muscle tissue of tamoxifen-injected chimeras (+) or non-injected control (-). Only the edited shorter 146bp (red asterisk) product was detected in both groups. **(M)** Representative flow cytometry analysis showing the *Pax7-nGFP*$^{+}$ / ITGA$^{+}$ satellite cells. **(N)** Pax7-nGFP expression in myoblasts that have been derived from ITGA7$^{+}$satellite cells of the indicated animals and conditions. Scalebar, 100µm.

**Figure S3**

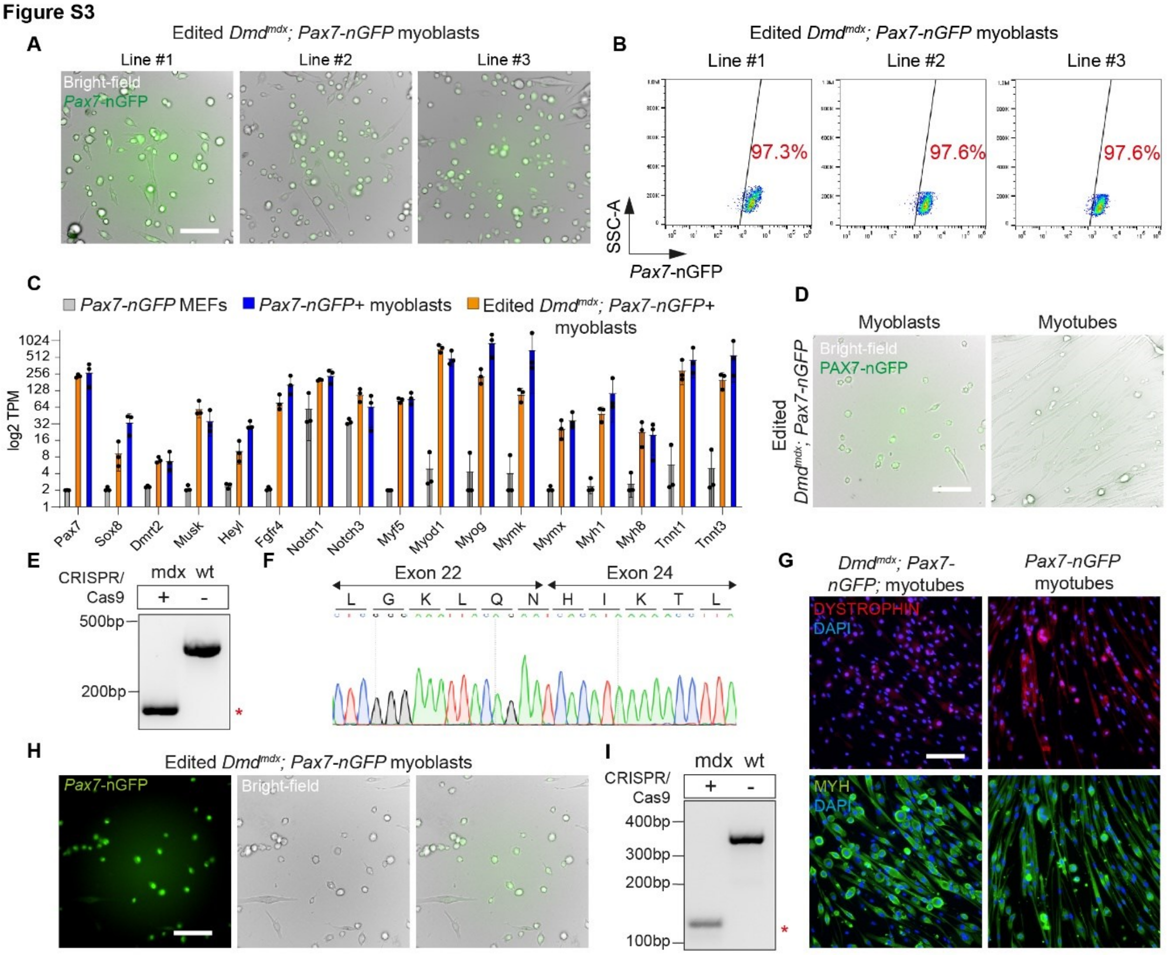


**Fig. S3.**

**(A)** Bright-field and microscopy images of myoblast lines isolated from three different intraspecies chimeras. Scalebar, 100µm. **(B)** Flow cytometry analysis for Pax7-nGFP in the indicated myoblasts lines. **(C)** Bar plots showing the log2 normalized expression of the indicated genes for edited *Dmd$^{mdx}$; Pax7-nGFP* myoblasts compared to *Pax7-nGFP* myoblasts and MEFs. N=3 cell lines per group. TPM, Transcripts per kilobase Million, MEFs, mouse embryonic fibroblasts. **(D)** Representative bright-field images showing edited *Dmd$^{mdx}$; Pax7-nGFP* myoblasts and myotubes. Note that GFP is downregulated upon differentiation. Scalebar, 100µm. **(E)** PCR for *Dystrophin* amplified using cDNA of edited *Dmd$^{mdx}$; Pax7-nGFP* myotubes and wt myotubes. Non-edited and edited *Dystrophin* are 396bp and 183bp (red asterisk), respectively. **(F)** DNA sequencing of (E) showing successful ligation of exon 22 and exon 24. **(G)** Immunofluorescence for dystrophin in the indicated cell lines. Scalebar, 100µm. **(H)** Bright-field and microscopy images of edited *Dmd$^{mdx}$; Pax7-nGFP* myoblasts prior to transplantation. Scale bar, 100µm. **(I)** PCR for *Dystrophin* in *Dmd$^{mdx}$; Pax7-nGFP* myoblasts showing only the edited shorter PCR product (red asterisk) vs. control.

Figure S4

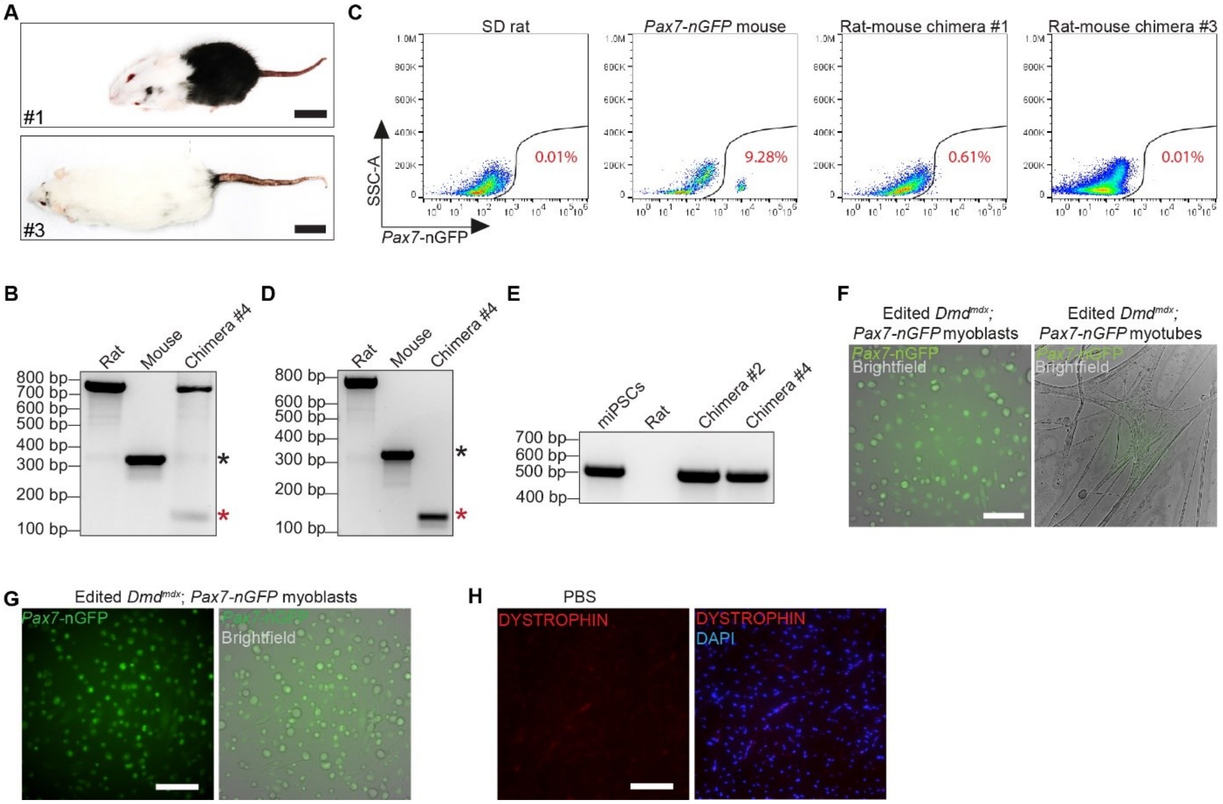


**Fig. S4.**

**(A)** Photos of rat-mouse chimeras #1 and #3 at 5 (top, chimera #1) and 17 (bottom, chimera #3) weeks of age. Scale bar, 3.5cm. **(B)** PCR for rat and mouse dystrophin from muscle lysates of the indicated animals. Black and red asterisks denote unedited (340bp) and edited (146bp) murine dystrophin, respectively. **(C)** Flow cytometry analysis of GFP expression in whole-body skeletal muscles isolated from the indicated animals. **(D)** PCR for rat and mouse dystrophin in myoblasts isolated from the indicated animals. Black and red asterisks denote unedited and edited murine dystrophin. **(E)** PCR for *Pax7-nGFP* transgene in myoblasts isolated from chimeras #2 and #4 and controls. **(F)** *Dmd*$^{mdx}$*; Pax7-nGFP* myoblasts and derivative myotubes. Scale bar, 100µm. Bright-field settings differed between images, whereas GFP channel was the same. **(G)** Edited *Dmd*$^{mdx}$; *Pax7*-nGFP myoblasts before transplantation. Scale bar, 100µm. **(H)** *Dmd*$^{mdx}$*-4Cv; Prkdc*$^{scid}$ TA muscle cross-section for PBS injected control. Scale bar, 100µm
